# Simplified Methodology for a Modular and Genetically Expanded Protein Synthesis in Cell-Free Systems

**DOI:** 10.1101/742726

**Authors:** Yonatan Chemla, Eden Ozer, Michael Shaferman, Ben Zaad, Rambabu Dandela, Lital Alfonta

## Abstract

Genetic code expansion, which enables the site-specific incorporation of unnatural amino acids into proteins, has emerged as a new and powerful tool for protein engineering. Currently, it is mainly utilized inside living cells for a myriad of applications. However, utilization of this technology in a cell-free, reconstituted platform has several advantages over living systems. The common limitations to the employment of these systems are the laborious and complex nature of its preparation and utilization. Herein, we describe a simplified method for the preparation of this system from *Escherichia coli* cells, which is specifically adapted for the expression of the components needed for cell-free genetic code expansion. In addition, we propose and demonstrate a modular approach to its utilization. By this approach, it is possible to prepare and store different extracts, harboring various translational components, and mix and match them as needed for more than four years retaining its high efficiency. We demonstrate this with the simultaneous incorporation of two different unnatural amino acids into a reporter protein. Finally, we demonstrate the advantage of cell-free systems over living cells for the incorporation of δ-thio-boc-lysine into ubiquitin by using the *methanosarcina mazei* wild-type pyrrolysyl tRNACUA and tRNA-synthetase pair, which can not be achieved in a living cell.

## INTRODUCTION

In nature, a native protein is limited to the 20 canonical amino acids and their specific chemical and physical properties. Genetic code expansion (henceforth GCE) expands this limit and offers the ability to site-specifically incorporate hundreds of new unnatural moieties by stop codon suppression. Thereby enabling the rational enhancement of the chemical and physical properties of proteins. This methodology was established in the early years of the current century in *E. coli*[1] and gradually was adapted to many other organisms in both eukarya[2–7] and prokarya[8,9].

Moreover, synthetic *E. coli* strains were generated to remove all UAG stop codons and their cognate release factor[10]. More recently, in a remarkable technical feat, the total synthesis of an *E. coli* genome was achieved. This was done specifically to afford a better application of this technology, freeing rare codons for the incorporation of unnatural amino acids (Uaas) [11]. The ability to produce genetically expanded proteins inside living cells resulted in many applications driven by hundreds of Uaas [12]. However, some of these amino acids are complicated to synthesize and expensive, thereby limiting its use in the relatively large volumes of the bacterial growth media. These amino acids have low permeability and sometimes are toxic to the host organism, thereby limiting their use in living cells. Cell-free protein synthesis with GCE capabilities alleviates these specific limitations. Another critical advantage for genetically expanded cell-free protein synthesis is the absence of the host genome, which removes limitations of toxicity and off-target suppression that may occur by suppression of the host endogenous stop codons. This is even more pronounced when two different Uaas are utilized at the same time; an exploit that is possible in living cells but is simplified in cell-free platforms[13].

The first genetically expanded cell-free protein synthesis system was, in fact, a precursor for the GCE systems in living cells, it was realized in the late 1980s by Peter Schultz’s team, using yeast Phe-tRNAs which were mutated to suppress the UAG stop codon in *E. coli* and was chemically aminoacylated by a Uaa. These tRNAs were added to *E*. *coli* cell extracts and successfully suppressed the designated stop codons[14]. After the successful *in-vivo* adaptation of GCE in the early 2000s, again by the Schultz team[15], the field has mostly moved to live cells as chassis. At the same time, Cell-free transcription-translation methodologies advanced to achieve higher yields[16,17], simplified and improved preparation protocols[18,19], and better understanding and control of the methodology itself[20]. As a result, GCE in cell-free protein synthesis also improved. The PURE system was introduced and achieved ca. 80 mg/mL yields of Uaa containing proteins[21]. The first systems which utilized both orthogonal tRNA and synthetase, exogenously added to *E. coli* extracts, were introduced by the Nishikawa group[22] without reported yields but with 50% suppression efficiency, and by the Swartz group, which also improved its yields and its suppression efficiency[23]. This methodology was further enhanced in recent years by the Jewett team, which implemented it in RF1-deficient cells[24,25]. However, the main problem of an approach where the orthogonal tRNA and synthetase are added exogenously is that it requires the purification of both components, a step which is both laborious and might be challenging with insoluble synthetases, as in the case of the pyrrolysyl tRNA synthetase.

To overcome this limitation, another approach has been developed, by both the Bundy team and by us, where the orthogonal tRNA and synthetase are transformed and expressed endogenously in the host cell, prior to extract preparation[26,27]. This approach completely circumnavigates the otherwise needed purification steps and achieves reasonable yields of up to 300 mg/mL with high suppression efficiency. As we recognize that the complex and laborious nature of the preparation and use of GCE in cell-free protein synthesis has been a barrier to the more widespread use of this methodology, this study will expound the extract preparation method of this approach in a straightforward manner. Moreover, we will present an adaptation of the protocol, which reduces its complexity and laboriousness, without compromising its quality. Lastly, we will promote a modular approach that was recently introduced[13], where different endogenous components could be expressed in separate bacterial extracts, stored for long periods and combined in a desired combinatorial fashion when needed [Fig 1].

**Figure 1.**
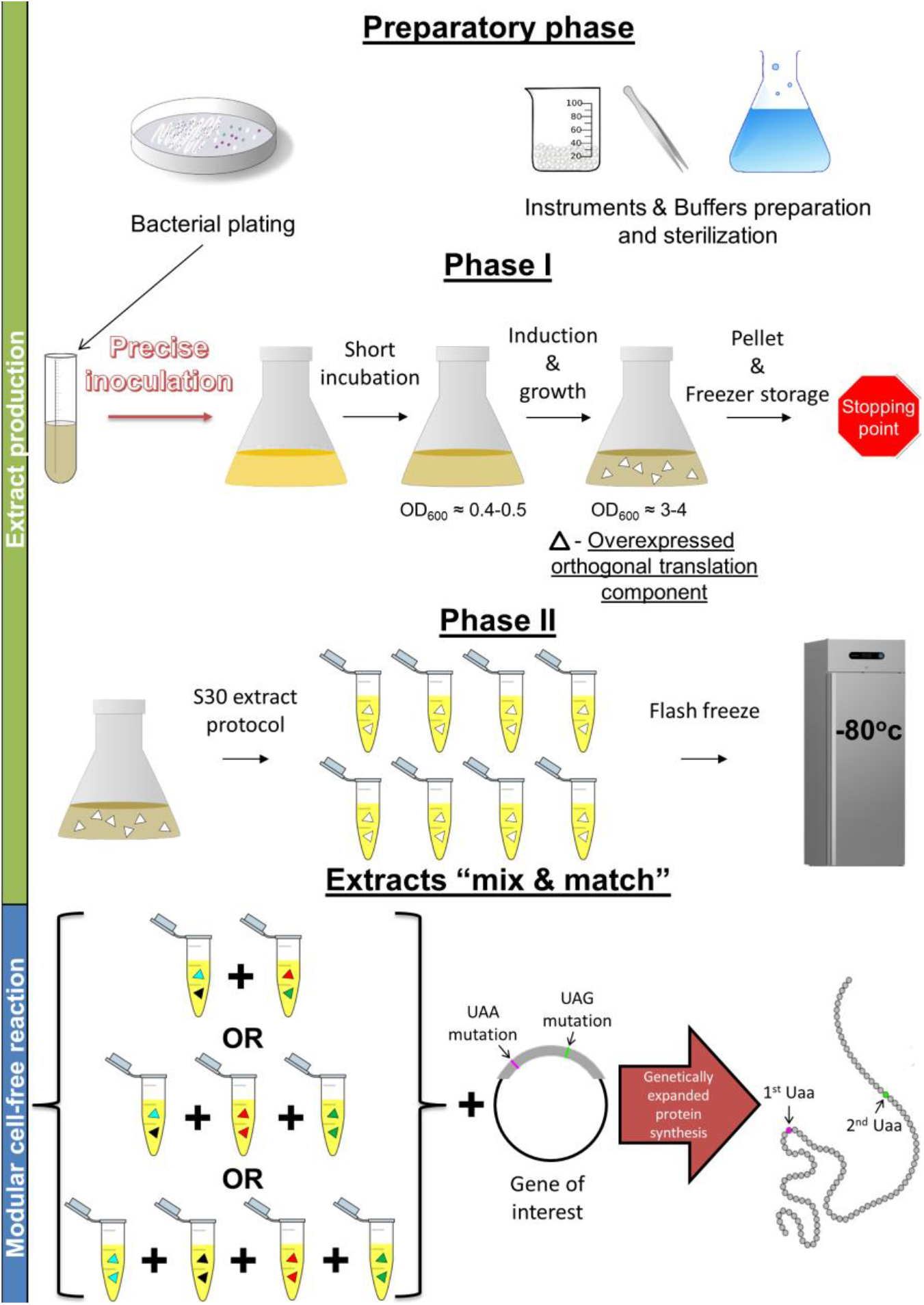
General scheme of the simplified preparation protocol and the modular approach for the cell-free reaction presented in this study. The entire protocol is divided into three stand-alone phases: i) the preparatory phase, where all necessary materials, instruments, solutions, and media along with initial plating of the specific *E. coli* strain are prepared. ii) Phase one: bacterial growth is scaled up and induced to produce the desired components. Next, the cells are harvested, pelleted, and stored in the freezer. iii) Phase two, the extract is prepared under the S30 protocol *(vide infra)* with small variations. Once several extracts are made, each containing different translational component pairs, the cell-free protein synthesis can be performed, where the modular reaction can produce one or more products while varying the components simply by mixing and matching extracts.

Herein, we present the utilization of this approach to simultaneously incorporate two different Uaas to the same protein. Finally, we demonstrate a new use of this approach, and cell-free GCE in general, by the incorporation of δ-thio-N-boc-lysine (TBK, Fig 2a) to reporter proteins and yeast ubiquitin, which is both challenging and expensive to accomplish inside living cells. We further demonstrate the ability to use this Uaa as a bio-orthogonal chemical handle for future native chemical ligation of ubiquitin and polyubiquitins.

**Figure 2.**
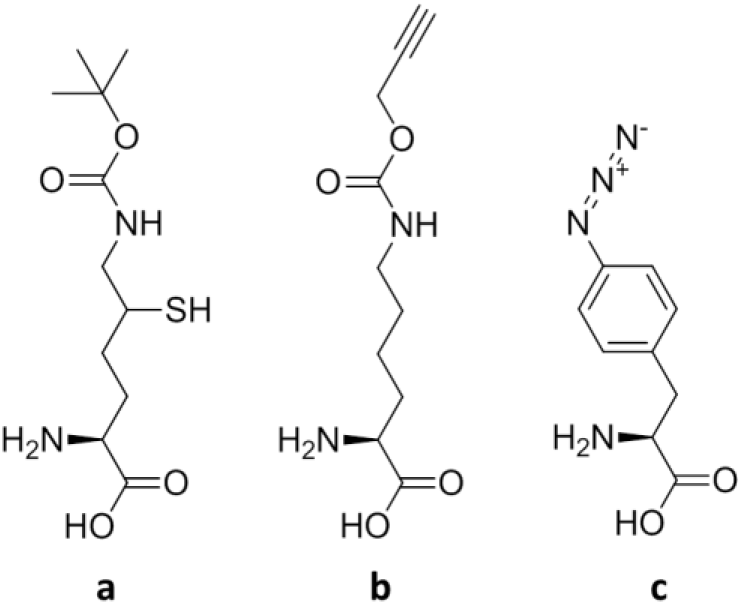
Chemical structures of Uaas used in the presented research. **a)** δ-thio-N-boc-lysine (TBK). **b)** N-propargyl-L-lysine (PrK). c) p-azido-L-phenylalanine (AzF)

## RESULTS AND DISCUSSION

### Simplified extract preparation and cell-free reaction for genetically expanded protein synthesis

Bacterial extract preparation is the most laborious phase in cell-free protein synthesis; if GCE is not necessary, it could be purchased commercially. However, if GCE is required, the extract must be prepared by the user as the orthogonal tRNA synthetase, and a tRNA pair are commercially unavailable. As we recognize that this phase is a major limitation for the employment of this approach, the first goal of this study is to simplify the process and improve its accessibility to potential users.

One of the most widely used protocols for extract preparation is the S30 protocol[13,18,24,27–32], which was modified in this study to create the genetically expanded cell-free protein synthesis system. When using orthogonal translation systems (OTS) to achieve GCE harbored on a replicating bacterial vector, such as the pEVOL[33] vector, the extract preparation protocol becomes much longer and more difficult as the bacterial growth becomes slower. Thus an induction step becomes necessary. The original S30 protocol starts with bacterial plating, it then requires three consecutive inoculation and growth rounds. In each round the culture volume is increased, and the rounds should be precisely timed to re-inoculate the bacteria in the late-log-phase. This leads to difficulties in the timing of inoculations when dealing with bacteria with slow growth rates. For example, the strain which was used in this study; *E. coli* C321ΔA[10] transformed with the pEVOL Pyl OTS vector, harboring the *Methanosarcina mazei (Mm)* pyrrolysyl orthogonal pair (henceforward Pyl-OTS). This strain has an average doubling time of ca. 50 minutes, which leads to difficulties in following the original protocol and its required consecutive inoculations. This problem is enhanced when induction is required, as it further reduces growth rates and adds another lag phase caused by the addition of the inducer. To address this problem, we have divided the protocol into three stand-alone phases which could be performed separately: The first, preparatory phase, which will not be elaborated as it follows the S30 preparations phase without significant alterations which were detailed by Sun et. al.[34]

#### Phase 1

In this sequential phase, bacterial scale-up, induction, protein overexpression, and preparation for lysis are performed. For this phase to fit in a standard workday, the S30 protocol had to be adjusted. We designed four different protocol groups with variation in the inoculation method: (1) Prepared under the original S30 protocol[27], with its three required consecutive inoculations. (2) The first and second inoculations were removed, and the final inoculation (to 1 L of growth media) was done directly from the plates, with a very low initial bacterial concentrations which then were incubated for overnight. (3) The second inoculation was removed, meaning that the bacteria were first inoculated from plates to 50mL of growth media and incubated for overnight, in which it reached stationary phase (instead of the required log phase). The final inoculation was done by transferring 10 mL of stationary-phase bacteria to 1 L of growth media (this volume corresponds to a 1/100 dilution, the dilution factor required by the original protocol). (4) Similar to 3, but the final inoculation was done by transferring 40 mL of stationary-phase bacteria (a dilution factor of 1/25). Using larger bacterial inoculation was done to obtain near-induction O.D._600_ (ca. 0.5), which allows induction to be performed after a brief (ca. 1 generation time) incubation period.

The three altered inoculation protocols all represent significant improvement as they enable the induction of the plasmid to be performed early in the workday (when the final culture reach O.D._600_ of ~0.5). Even if induction is not needed, these altered protocols still sidestep timing limitations and reduce the duration of inoculation time from three days to only two days. After induction, final growth and OTS expression takes until around noon. In this step, we have found that the requirement in the original S30 protocol of cell harvest when cultures reach O.D._600_ of 1. 5-2 could be increased to 3-4 thereby doubling the biomass and the resulting extract without any apparent compromise of efficiency in the system [Fig 3b]. Lastly, the regular washing step is conducted in the afternoon

**Figure 3.**
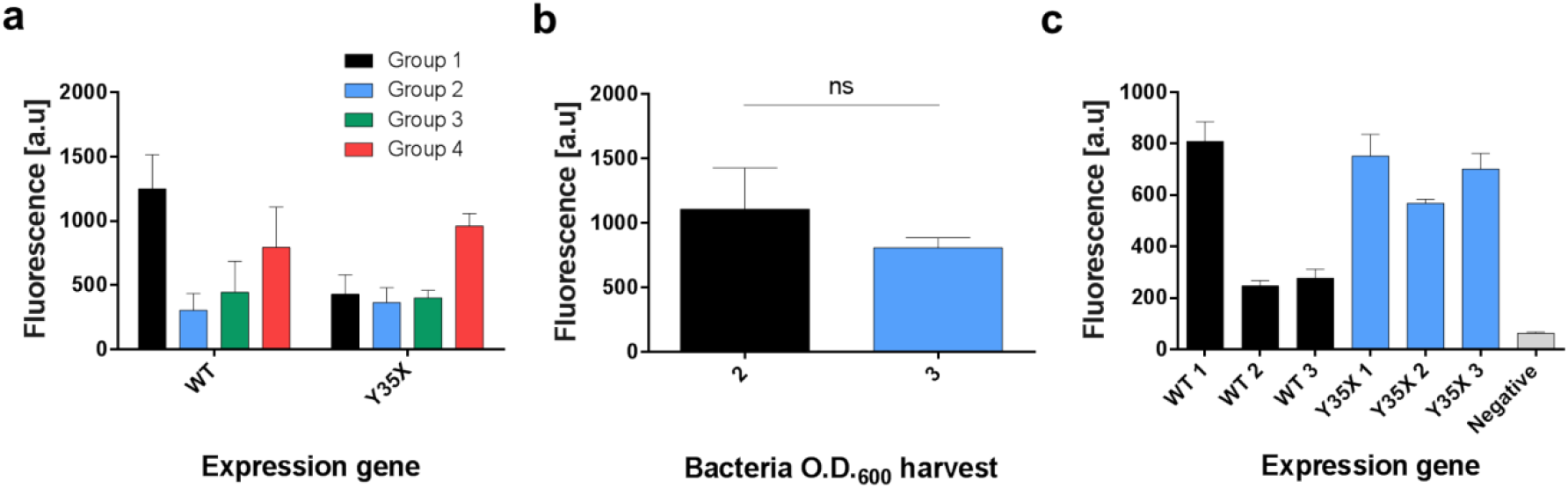
Comparison of performance using different extract preparation protocols. **a)** Cell-free production of WT GFP and genetically expanded Y35PrK GFP. Fluorescence intensity of produced WT and mutant GFP for four protocols: 1) S30 protocol 2) No intermediate inoculation, log phase maintained 3) No intermediate inoculation, stationary phase instead of log phase added to final culture in small volume 4) Similar to 3, but added in large volume to final culture to obtain near-induction O.D. **b)** Cell-free production of WT GFP with extracts harvested at different O.D._600_. Original S30 protocol suggested bacterial harvest at O.D.≈ 2, compared to higher bacterial harvest at O.D.≈ 3. The results are not significantly different (two sided T test, t=1.57, df=4, p-val=0.19). **c)** Cell-free production of GFP for comparison between different plasmids, using Pyl-OTS for UAG suppression. Fluorescence intensity of expressed protein from three different batches of WT GFP and Y35X plasmids. Each plasmid type has the same sequence but was purified independently.

Next, we set out to find a stopping point for the extract preparation protocol, which is not mentioned in the available S30 protocols. We have reasoned that it could be stopped after the washing phase, for a prolonged period, as a bacterial pellet before lysis. All of the groups mentioned above, excluding group 1, were subjected to this stopping point. This roughly divided the protocol midway, which spreads the intensive work of one day to two more convenient days of work. We have tested the resulting extracts from all protocol combinations with the cell-free expression of wildtype (WT) GFP and a GFP gene mutated at position 35 from tyrosine (UAC) to amber stop codon (UAG), labeled as GFP Y35X. The reaction mixture of GFP Y35X was supplemented with 1 mM of the Uaa N-propargyl lysine (PrK, Fig 2b) which could be incorporated by UAG suppression by the Pyl-OTS [Fig 3a]. For the GFP Y35X protein, we found that the new protocols including the inoculation and the stopping point alterations do not compromise the yields nor the fidelity of the protein with an incorporated Uaa, while significantly reducing the workload [Fig 3a]. It appears, however, that group 4 attains the best results under these specific conditions.

#### Phase 2

In this phase, the rest of the S30 protocol is performed including lysis (using a bead-beater), S30 separation by centrifugation of lysate, nuclease digestion by incubation and dialysis. In this phase we have found two notable adjustments, i) we have found that the dialysis should not be continued for more than 3-4 hours in 4^0^C as the efficiency of the genetically expanded system decreases after prolonged dialysis, presumably due to the lower stability of the orthogonal tRNA. And ii) The final extract protein concentration measurement and calibration steps, in the original protocol, are both time-consuming and do not significantly affect the efficiency of the extract. Therefore, this phase should be considered optional and should be applied only when the complete optimization of the extract is needed.

#### Cell-free transcription/translation reaction

The genetically expanded protein synthesis reaction requires further consideration. First, the GCE cell-free buffer calibration is an important step which effects the yields of protein synthesis. In some S30 protocols, it is advised to calibrate the Ion concentrations of K-glutamate and Mg-glutamate and the concentration of DTT. However, in genetically expanded cell-free protein synthesis, we have found that concentrations usually could be fixed to 100 mM and 2.5 mM for K-glutamate and Mg-glutamate, respectively without DTT, with no significant effect on the system yields. Second, the Uaa concentration was found to achieve the best yield in the concentration range of between 0.5 mM (for N-Boc-Lysine)[27] and up to 3.6 mM (for TBK, in this study), an optimal concentration should be calibrated for each specific Uaa while the starting concentration should be 1 mM. Finally, we note that cell-free reaction yield-variability is relatively high; this warrants separate attention and further investigations as to its sources. However, it is important to discuss two sources of this variability, which could be a common impediment in the employment of this system. First, the context and location of the site chosen and mutated to facilitate Uaa incorporation have a significant effect on the efficiency of the system[35,36], but notably, from our experience, the context, and locations effects could be different between *in-vivo* and cell-free systems [data not shown]. This subject warrants a systematic investigation which is outside the scope of this study. However, it will be wise to screen several sites of the protein to achieve successful cell-free Uaa incorporation. And second, the expression plasmid, which is added to the reaction mixture carries significant variability. We have noticed, on numerous occasions, that batches of the same plasmid, from unknown reasons, could vary considerably when added to reactions and in extreme cases, there could be no expression at all. To demonstrate this point, we have tested three batches of two plasmids, pBEST GFP WT, and pBEST GFP Y35X, in an otherwise identical reaction conditions [Fig 3c]. The results indicate the extent of the variability that arises solely from differences in plasmid batches. Therefore, it is beneficial to test several plasmid batches, sometimes even by using different prep kits, before discarding an extract batch. The source of this variability is unknown to us at this point, and it warrants further attention.

### A modular approach to cell-free protein synthesis with Uaas

Recently, we have found that extracts from the same bacterial strain but with different genotypes and from different batches could be mixed and combined[13], this led us to develop a modular approach for cell-free protein synthesis with GCE. In this approach, a component of choice, which could be a protein or a non-protein-coding gene like tRNAs, could be expressed separately in bacteria which will sequentially be prepared as an extract that contains the said component. These separate extracts could be stored for an extended period of time (some extracts used in this study were stored for over four years), thawed upon demand and be combined, mixed and matched according to the experimental design. Specifically, we have utilized this approach to express a reporter protein containing two different Uaas. A GFP gene was mutated in position 193 to UAG and position 35 to UAA: “double mutant.” We have tested the GFP expression in two modular combinations with and without the supplementation of the Uaas as a negative control [Fig 4a]: 1) a two-extracts system, one which contains the Pyl-OTS which incorporate PrK in response to UAG codon, the other contains the p-azido-L-phenylalanine (AzF)-OTS which incorporate AzF in response to UAA codon. Both OTSs contain a tRNA synthetase and a tRNA pair of genes. 2) a four-extracts system each contains only one of the four above mentioned components: 1) AzF synthetase 2) AzF tRNA for UAA suppression 3) PrK synthetase 4) PrK tRNA for UAG suppression. The results clearly show that this approach is viable for at least four different combinations of interacting components of tRNAs and tRNA synthetases. However, there is a noticeable decrease in the system yield between the 2-extracts system and the 4-extracts system. We reasoned that this decrease in yield is a result of the relative dilution of each component in the final volume, as more extracts are mixed. To test this, we have diluted the AzF-OTS with increasing concentrations of Pyl-OTS and quantified the double mutant production rates [Fig 4b]. The results show a significant decrease in the expression level of the double mutant, while only a slight decrease is observed in the level of WT protein, which does not require increasingly diluted components. We have fitted the results with linear regression [Fig 4c], so it explains ca. 97% of the variability in proteins levels that we observe, the two-extracts, and the four-extracts modular systems results are in agreement with the linear model and fall within the standard deviation of the model.

**Figure 4.**
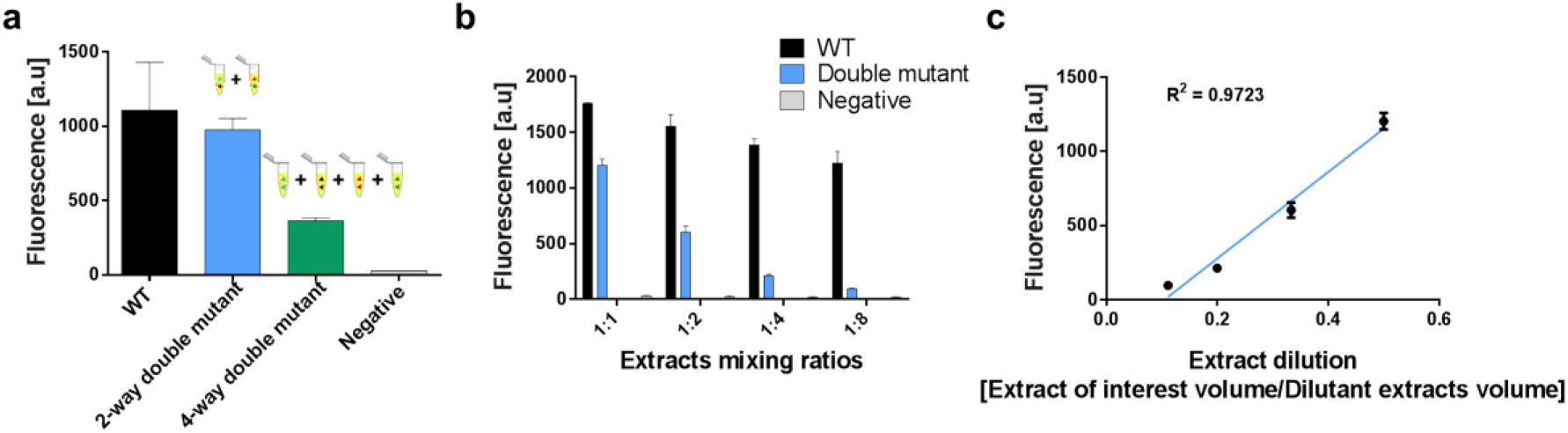
Cell-free production of double mutant GFP protein (Y35UAAD193UAG), using extracts mixtures of different ratios. Fluorescence intensity of double mutant production was compared to WT protein or double mutant without any Uaas (negative control). **a)** mixtures of OTS pairs and mixtures of OTS single components extract comparison for double mutant GFP production. A two-extracts mixture is composed of AzF-OTS for UAA suppression and Pyl-OTS for UAG suppression, while a four-extracts mixture composed from the following extracts: 1) AzF synthetase 2) AzF tRNA for UAA suppression 3) PrK synthetase 4) PrK tRNA for UAG suppression. **b)** OTS pairs extract mixture, between AzF-OTS for UAA suppression and increasing quantities of Pyl-OTS for UAG suppression, at different ratios. **c)** Linear regression of extract dilution for double mutant GFP expression measured by GFP fluorescence.

### Incorporation of δ-thio-N-boc-lysine (TBK) into reporter proteins and Ubiquitin

Synthesis of the Uaa, TBK was performed using a published procedure adapted from Virdee et al. [37]. The TBK amino acid is being used for bio-orthogonal chemistry of native chemical ligation, which results in an isopeptide bond formation between two proteins of interest. This Uaa was shown to be incorporated exclusively *in-vivo* by the pyrrolysyl orthogonal translation system with an evolved pyrrolysyl tRNA synthetase[37]. In our experience, the *in-vivo* utilization of this Uaa was both expensive (as high concentrations in addition to large culture volumes were required) and achieved low yields, which were insufficient for downstream applications. Therefore, TBK incorporation was tested in the genetically expanded cell-free protein synthesis using the modular extract with the Pyl-OTS. Using this system, the TBK was first incorporated into GFP in position 35 (i.e., GFP Y35X) and RFP in position 15 (i.e., RFP D15X). The incorporation of the Uaa was tested using kinetic fluorescent measurements for both GFP [Fig 5a] and RFP [Fig 5b].

**Figure 5.**
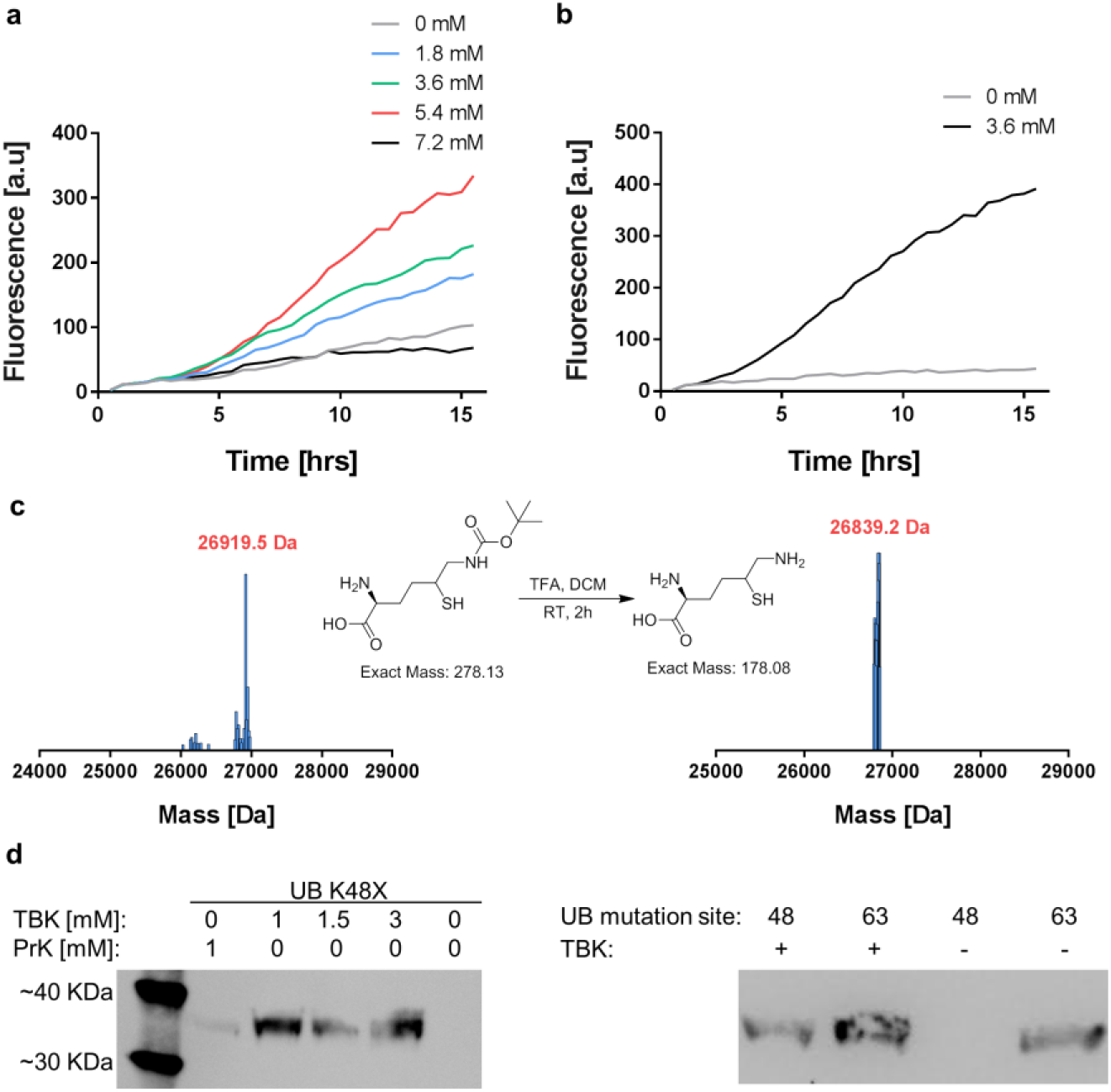
TBK incorporation into proteins, using genetically expanded cell-free protein synthesis. **a)** Cell-free kinetics of GFP Y35X production in the presence of different TBK concentrations. **b)** Cell-free kinetics of RFP D15X production in the presence and absence of TBK. **c)** ESI mass spectra of purified RFP D15X with TBK, before and after boc-deprotection. **d)** Anti-ubiquitin western blot of genetically expanded cell-free protein synthesis of the 36.4 kDa Ub-Intein-CBD construct. On the left panel, Cell-free reactions to incorporate PrK and TBK in different concentrations. The Ub K48X without Uaa lane serves as a negative control. On the right panel, Comparison between TBK incorporated into two different branching sites, K48X and K63X in the Ubiquitin construct in the presence or absence of TBK, the band in site 63 in the absence of TBK could be a result of stop codon readthrough.

To be utilized for native chemical ligation, the TBK must first be boc-deprotected. To test this, we have expressed the RFP D15TBK variant in a large volume, purified it using nickel affinity chromatography and removed the protective group using TFA, and the resulting protein RFP D15TK was validated using mass spectrometry and compared to the original compound (Fig 5c).

To demonstrate the specific advantage of cell-free protein synthesis, we have sought to incorporate TBK to the ubiquitin gene. Ubiquitin is a highly conserved 76 amino acids protein that is present in all the eukarya which serve as a post-translational modification of other proteins. This system was discovered in the context of protein degradation[38] since it was found to play a role in a myriad of cellular processes. Every ubiquitin can be ubiquitinated by one or more other ubiquitins at seven different sites, thus creating a complex code[39]. Efficient methods for the synthetic generation of this code is a feat which is long sought after. Therefore, we have decided to take a step in this direction. The yeast *UBI4* gene (the ubiquitin gene) was amplified from the yeast genome and fused to intein-chitin binding domain (Int-CBD) construct. This construct enables the formation of N-terminal-thioester and affinity purification[40]. The 36.4 kDa construct was cloned to the pBEST plasmid to be expressed in the genetically expanded cell-free protein synthesis system. Its expression was tested to incorporate both PrK or varying concentrations of TBK at the poly-ubiquitin branching lysine site K48 [Fig 5d, left panel] and was also validated for a second branching site – K63 [Fig 5d, right panel]. These produced moieties represent a first step towards the synthetic synthesis of a branched poly-ubiquitin code that could be generated using native chemical ligation. This is achieved as this ubiquitin construct can act both to ubiquitinate utilizing intein cleavage to create an N-terminal thioester or be ubiquitinated utilizing the boc-deprotected moiety of the TBK Uaa in a combinatorial manner.

## Conclusions

Herein we present a simplified method to generate genetically expanded cell-free protein synthesis system. It reduces the laborious nature of the protocol by altering the inoculation and adding possible stop points. Also, this protocol reduces the complexity of the S30 protocol by removing optional steps such as extract calibrations. Once a simplified protocol for extract preparation is used, it could be utilized for several genotypes of the same bacterial strain, each over-expressing a different set of components. These different genotypic extracts can contain various components which could be stored for a long period while maintaining most of its functionality and then mixed-and-matched in any combinatorial way without the need for a new preparation. This results in a modular approach to cell-free protein synthesis with genetic code expansion. We anticipate that these results will increase the use of genetically expanded cell-free protein synthesis, in the specific cases where it presents an advantage over living cells. We acknowledge that the modular approach presented herein is not limited only to GCE, but it could directly be adapted and utilized for other applications. These applications include chaperon-required protein expression, an amalgamation of toxic or aggregating components which cannot be co-expressed in living cells, and any cell-free protein expression which requires several components which it would be beneficial to combinatorically mix and match between them.

## METHODS

### Strains and plasmids

The bacterial strain used for the preparation of all cell extracts is the C321Δ*prfA* strain (Addgene #48998) carrying a pEVOL plasmid, transformed before growth and extract preparation. In this paper, six types of pEVOL plasmids were used: 1) *Methanosarcina mazei (Mm)* orthogonal pair of *Mm*-PrKRS\*Mm*-tRNA_CUA_^PrK^ (Pyl-OTS) 2) *Methanocaldococcus jannaschii (Mj)* orthogonal pair of *Mj*-AzFRS\*Mj*-tRNA_UUA_^AzF^ (AzF-OTS) 3) *Mm*-PrKRS (no tRNA) 4) *Mm*-tRNA_CUA_^PrK^ (no synthetase) 5) *Mj*-AzFRS (no tRNA) 6) *Mj*-tRNA_UUA_^AzF^ (no synthetase)[13].

All expression genes targeted for CFPS were harbored on a pBEST plasmid[41]. pBEST deGFP[27] was expressed in three forms, WT GFP, Y35UAG GFP, Y35UAA D193UAG GFP. All mutations were introduced using standard mutagenesis protocol.

The pBEST-UB constructs were generated as follows. The *S. cerevisiae UBI4* ubiquitin gene was amplified from its genome. Next, it was fused, in its N terminus to intein and CBD genes from the IMPACT kit [New England Biolabs] and the entire construct was amplified and cloned to the pBEST plasmid. pBEST UB was expressed in three forms, WT UB, K48UAG UB, K63UAG UB. All mutations were introduced using standard mutagenesis protocol.

### Extract preparation

In general, we have followed the Sun et al. visualized protocol[34]. The following are the specific changes made in this study. C321.Δ*prfA* bacteria containing a pEVOL plasmid were grown in four different conditions prior to lysis: 1) Prepared according to the original protocol[34] as control. 2) Bacteria were plated overnight. Next, an isolated colony was inoculated directly into the final culture in a 5 L Erlenmeyer with 1 L of growth media (the exact composition of the growth media is detailed in the original protocol). Meaning that the first and second inoculation steps from the original protocol were removed. The final culture was incubated for 12 hours after which its O.D._600_ was measured to be 0.14. Note that ideally, the user should calibrate the exact incubation time to reach an induction O.D._600_ of ~0.5 if using this condition. 3) Bacteria were plated for overnight. Next, an isolated colony was inoculated directly to the second culture in a 250mL Erlenmeyer with 50mL of growth media. Meaning that the first inoculation step was removed. The final inoculation was done after overnight incubation of the previous inoculation using 10 mL of deep-stationary phase bacteria. 4) Similar to 3, but added 40mL instead of 10mL of stationary-phase bacterial culture. This was done to obtain near-induction O.D. For all protocol groups, Induction of the pEVOL plasmid (Ara promoter), was done using 0.5% of L-arabinose when culture O.D._600_ reached 0.5. Following induction, all four protocol groups were incubated, and the bacterial culture was allowed growth and induced protein expression up to O.D._600_ of 3-4 (higher than the original protocol), in all four experimental conditions, the biomass was collected, dried, weighed, and stored in the −80°c. This is a stopping point we have added to the original protocol. In the following day (not a requirement, could be stored for longer periods), the biomass underwent the S30 cell extract protocol following the original prorocol[34] and resulted with a CFPS extract for genetic code expansion. The final extract was aliquoted into Eppendorf tubes in 30 μL batches, without any protein concentration measurement or calibration (in contrast to the original protocol).

### Cell-free protein synthesis reaction

The CFPS reaction volume was carried in a Nunc (black, flat transparent bottom) 384 well plate (Thermo scientific) in an incubated Synergy HT plate reader (Biotek), in temperature (29°c) for ~17 hours with intervals of 30 minutes. GFP expression was measured with an excitation wavelength of 488 nm and an emission wavelength of 507 nm.

In each cell-free reaction, 33% of the final reaction volume was composed of bacteria extract. When a single type of extract was used, all 33% were composed of that same extract. However, when several types of extracts were used, 33% of reaction volume was being divided by the different extracts. Different extracts ratios were tested when combining two separate extracts (1:1, 1:2, 1:4, 1:8 – ratio between extract of interest volume/dilutant extracts volume) and up to four separate extracts were combined at a 1:1:1:1 ratio.

In each cell-free reaction, 41.6% of the final reaction volume was composed of reaction buffer. The reaction buffer consists of the following compounds (concentrations are in final reaction values): 40.19 mM HEPES pH 8,1.21 mM ATP and GTP, 0.72 mM CTP and UTP, 0.16 mg/mL tRNA, 0.21 mM coenzyme A, 0.27 mM NAD, 0.60 mM cAMP, 0.055 mM folinic acid, 0.80 mM spermidine, 24.11 mM 3-phosphoglyceric acid,1.21 mM each of 20 amino acids, 1.61% PEG-8000, 96.47 mM K-glutamate, 2.41 mM Mg-glutamate (reagents were purchased from the same vendors as listed in Sun et al. 2013[34]. The buffer amount per reaction and the expression plasmid concentration were performed as described before[13]. All plasmids were purified from *E. coli* DH5a cells using the Wizard Plus SV minipreps kit [Promega].

Protein expression of genetically expanded proteins with stop codons mutations was done in the presence of 1 mM final concentration per Uaa, unless stated otherwise.

### UB western blot

Cell-free protein synthesis samples were diluted by a factor of ten and loaded into a 4-20% SDS gels [Genscript]. After transfer, anti-ubiquitin antibodies were used, the membrane was visualized using ImageQuant LAS 4000 imager [Fujifilm].

### Synthesis of δ-Thio-Boc-Lysine

The synthesis was performed as described by Virdee et. al.[37] and validated using mass-spectrometry (Finnigan Surveyor/LCQ Fleet, Thermo Scientific).

### Protein purification and mass-spectrometry analysis

Proteins were fused to 6xhistag and purified using standard nickel-column affinity purification. Purified protein samples were analyzed by LC-MS (Finnigan Surveyor/LCQ Fleet, Thermo Scientific).

## ACKNOWLEDGMENTS

We wish to thank the Azrieli (Y.C.) and Darom PhD fellowships (Y.C., E.O.) for supporting this study.We would like to greatfully acknowledge an ERC-StG grant number 260647 (L.A.) for supporting parts of the studies mentioned herein.

## AUTHOR CONTRIBUTIONS

Y.C and E.O share equal contribution to this paper, Y.C conceived and performed all experimentation, synthesized the molecules and co-authored the manuscript, E.O conceived and performed all experimentation and co-authored the manuscript, M.S. performed the TBK incorporation experiments, B.Z performed the extract preparation optimization experiments, R.D synthesized the TBK, L.A supervised, designed and conceived experimentations and manuscript preparation.

